# GenoTools: An Open-Source Python Package for Efficient Genotype Data Quality Control and Analysis

**DOI:** 10.1101/2024.03.26.586362

**Authors:** Dan Vitale, Mathew Koretsky, Nicole Kuznetsov, Samantha Hong, Jessica Martin, Mikayla James, Mary B Makarious, Hampton Leonard, Hirotaka Iwaki, Faraz Faghri, Cornelis Blauwendraat, Andrew B. Singleton, Yeajin Song, Kristin Levine, Ashwin Ashok Kumar Sreelatha, Zih-Hua Fang, Mike Nalls

## Abstract

GenoTools, a Python package, streamlines population genetics research by integrating ancestry estimation, quality control (QC), and genome-wide association studies (GWAS) capabilities into efficient pipelines. By tracking samples, variants, and quality-specific measures throughout fully customizable pipelines, users can easily manage genetics data for large and small studies. GenoTools’ “Ancestry” module renders highly accurate predictions, allowing for high-quality ancestry-specific studies, and enables custom ancestry model training and serialization, specified to the user’s genotyping or sequencing platform. As the genotype processing engine that powers several large initiatives, including the NIH’s Center for Alzheimer’s and Related Dementias (CARD) and the Global Parkinson’s Genetics Program (GP2). GenoTools was used to process and analyze the UK Biobank and major Alzheimer’s Disease (AD) and Parkinson’s Disease (PD) datasets with over 400,000 genotypes from arrays and 5000 sequences and has led to novel discoveries in diverse populations. It has provided replicable ancestry predictions, implemented rigorous QC, and conducted genetic ancestry-specific GWAS to identify systematic errors or biases through a single command. GenoTools is a customizable tool that enables users to efficiently analyze and scale genotype data with reproducible and scalable ancestry, QC, and GWAS pipelines.

## Introduction

Population genetics has experienced an explosion in sample sizes, fueled by advancements in microarray and sequencing technology paired with decreasing costs for compute. Along with the increase in the size of these studies comes an increase in the complexity of ensuring data quality and sample management. In this context, we present GenoTools, a Python package specifically designed to streamline and enhance the efficiency of genotype data processing. Population genetics studies demand a standard set of quality control measures to mitigate systematic biases that may affect downstream statistical analyses and machine learning. GenoTools addresses this critical need by integrating standard genotype data processing steps into one seamless, fully customizable pipeline that vastly simplifies the path from raw data to results and harmonizes datasets for federation and meta-analyses. The use of PLINK^1^ data types, a standard in population genetics, ensures compatibility and ease of data integration. GenoTools’ integration of array and whole genome sequencing (WGS) data genetic ancestry prediction, QC, and other comprehensive functionalities in a user-friendly design ensures researchers can easily build and scale custom pipelines to meet the specific needs of studies of any size from small studies to large biobanks.

## Methods

### Standard Pipeline

The standard GenoTools pipeline starts with genetic ancestry prediction. Samples are split by predicted ancestry and output to individual PLINK2 pgen format files, where they are run through detailed QC steps, default thresholds, and additional detail in the “**Quality Control**” section. Samples are evaluated for call rate, sex concordance, kinship, and heterozygosity. Variants are then excluded for non-random missingness by haplotype, Hardy-Weinberg equilibrium deviations in controls, and variant-level genotype missingness. Samples are output to the final PLINK2 pgen file type stratified by genetically ascertained ancestry group based on population references. Using the final output genotypes, principal components (PCs) are calculated using PLINK, and GWAS is run using the PLINK2 association module. Genomic Inflation (λ) factors are then calculated to check for QC/association quality. The full pipeline is shown in **Figure 1**.

**Figure 1.**
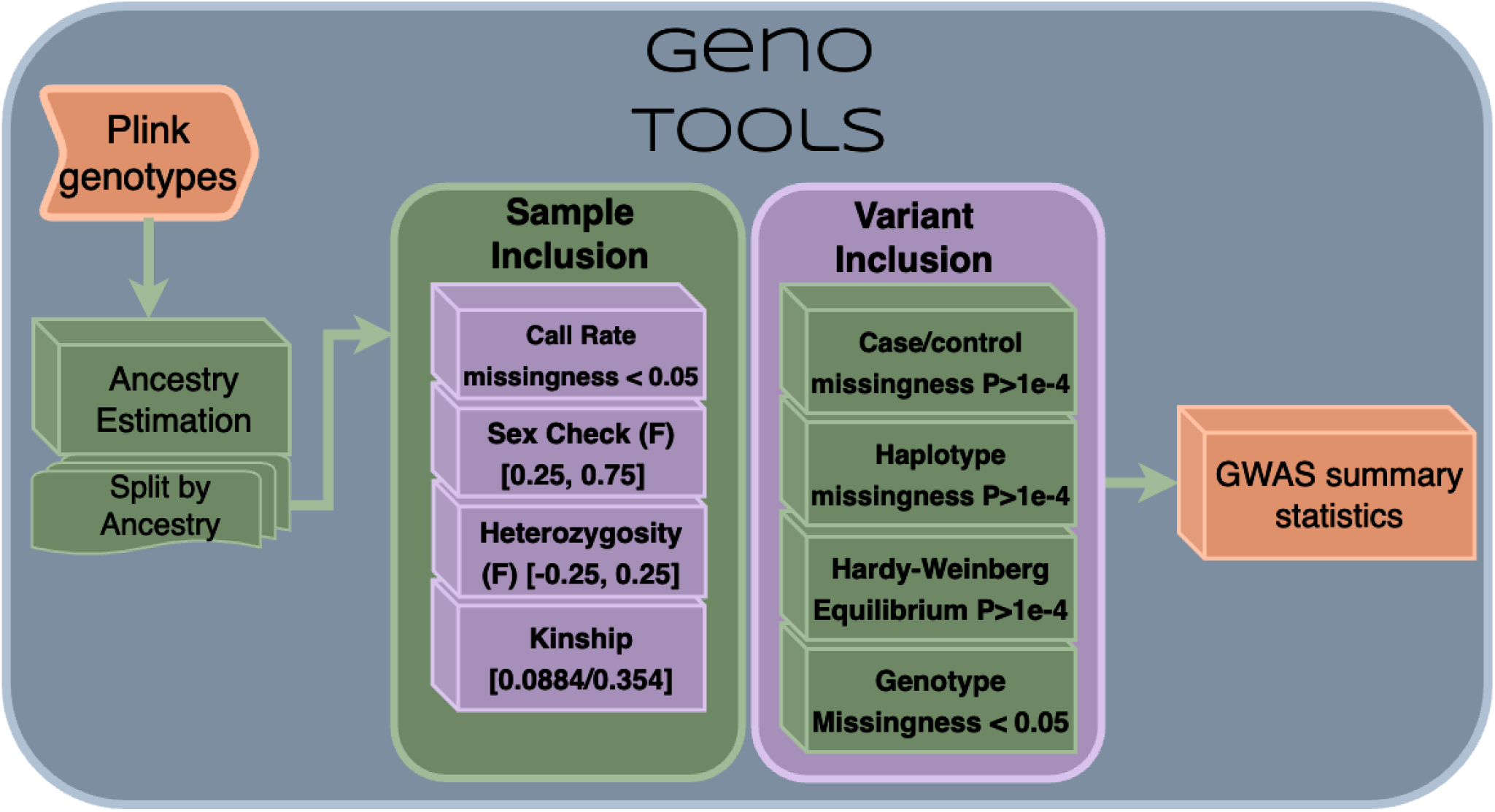
GenoTools Core pipeline

### Ancestry

GenoTools includes a suite of ancestry methods that includes ancestry-specific variant-level QC, custom model training, and prediction (**Figure 2**).

**Figure 2.**
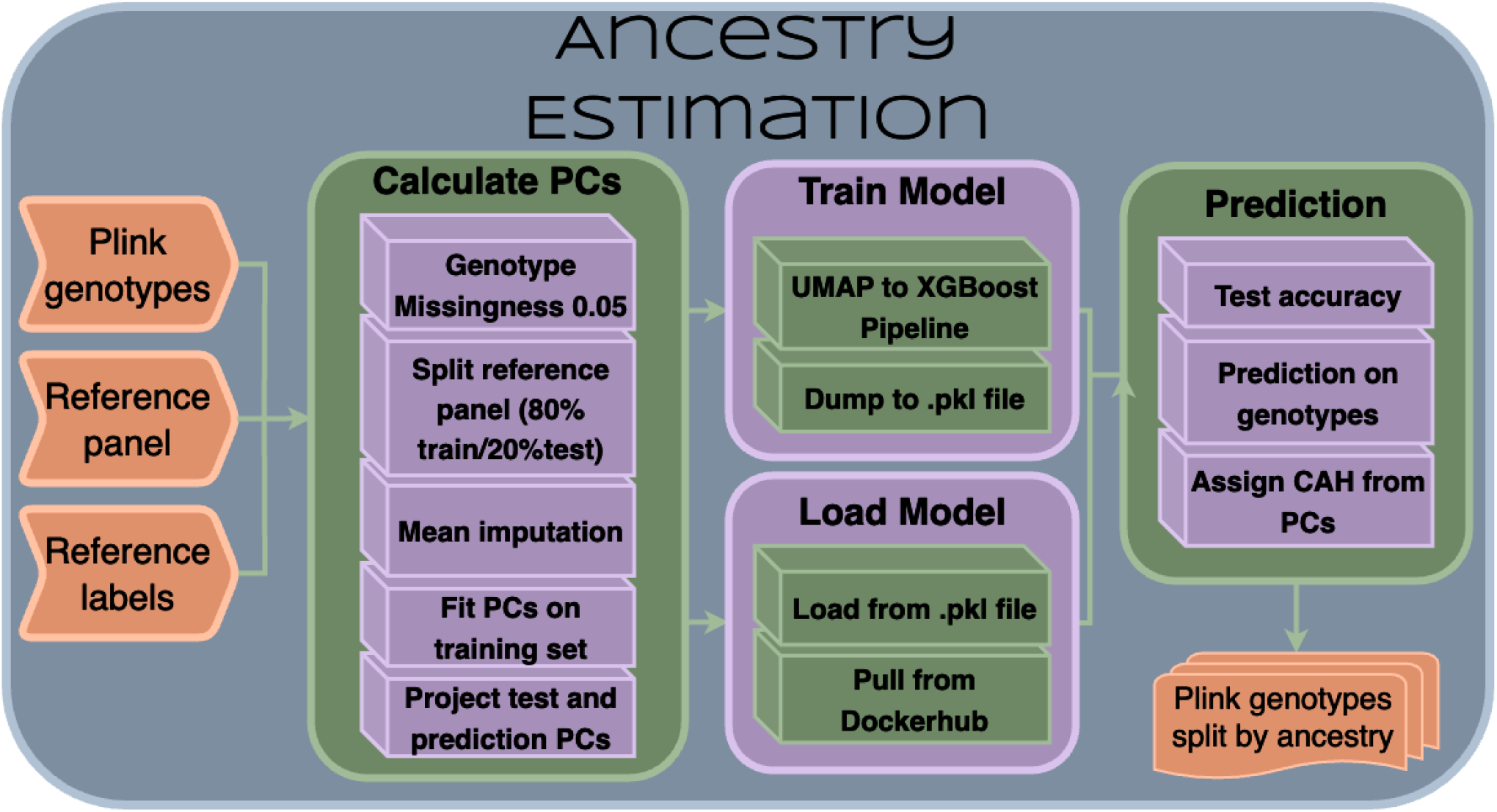
Ancestry Pipeline

### Model Training

The GenoTools ancestry pipeline begins by identifying overlapping variants between the reference panel and the prediction genotypes and matching the alleles to the reference panel. Reference samples are partitioned into a stratified 80/20 train/test split. 50 PCs are computed using PCA in scikit-learn^2^ after performing a FlashPCA-style normalization^3^. For the test and prediction sets, PCs are derived from the training set, fit, and scaled using the training set’s mean and standard deviation as previously described.

Subsequently, we integrate Uniform Manifold Approximation and Projection (UMAP)^4,5^ for further dimensionality reduction and Extreme Gradient Boosting (XGBoost)^6^ for classification. This step involves constructing a pipeline that combines UMAP and XGBoost Classifier with a gblinear booster.

Hyperparameter optimization is achieved through a grid search approach employing the following parameter grid:

- UMAP Parameters:
- Number of neighbors (umap n_neighbors): [5, 20]
- Number of components (umap n_components): [15, 25]
- Parameter ‘a’ (umap a): [0.75, 1.0, 1.5]
- Parameter ‘b’ (umap b): [0.25, 0.5, 0.75]
- XGBoost Parameters:
- Lambda (xgb lambda): Range from 10e−3 to 10e2 with steps in powers of 10.

Cross-validation within the pipeline is conducted using a five-fold StratifiedKFold strategy with random shuffling. Balanced accuracy is used as the scoring metric during the grid search optimization.

Upon model training, we extract and report the top results based on rank test score. We calculate and present the training balanced accuracy along with its 95% confidence interval. Similarly, the balanced accuracy on the test set and its 95% confidence interval are determined. During evaluation, a confusion matrix is generated based on test set predictions. Finally, the best-performing model from the grid search is serialized for predictions with Pickle in Python for future predictions and transfer learning applications.

### Ancestry Prediction

GenoTools enables the output of serialized models, tailored to specific genotyping arrays. This section outlines how these models render predictions on new datasets.

Overlapping SNPs between the user’s genotypes and the set of training variants are identified. Any variants that were used in model training that are not available in the new genotypes are imputed as homozygous reference. The reference samples are again partitioned into a stratified 80/20 train/test split, and the remaining missing prediction genotypes are imputed using the mean SNP value from the training data. PCs are computed and projected in the same manner as previously described. The projected PCs from the new samples are used as input for the serialized model, and, after being passed through the UMAP to the XGBoost pipeline, a list containing the predicted ancestry for each sample is returned as output. Please note that if the SNP overlap between the new samples and the variants used to train the selected model is insufficient, too many variants will be imputed as homozygous reference, and the rendered predictions will not be accurate. In this case, it is recommended to try another pre-trained model or to train a model that is specific to the new dataset, which can be done in GenoTools.

While the constructed reference panel does an adequate job of rendering accurate predictions for most samples, certain ancestry groups need to be better represented. For example, native South Africans, or Afrikaners, are known to have a complex genetic background with European, Asian, and African admixture^7^. Due to the lack of publicly available reference samples for populations such as Afrikaners and other highly admixed groups, GenoTools employs a method to identify samples of this nature and place them in an ancestry group that is not present in the reference panel, named “Complex Admixture History” (CAH). For population genetics, highly admixed samples of this nature should be analyzed independently of other ancestry groups.

Since there are no samples in the reference panel to base the prediction of CAH ancestry on, a PC-based approach is used instead. Using the training data, the PC centroid of each reference panel ancestry group is calculated, along with the overall PC centroid. The PC distance from each centroid is calculated for each new sample. Any sample whose PC distance is closer to the training data’s overall PC centroid than any reference panel ancestry group centroid is labeled as CAH. This method allows for samples to be labeled as CAH regardless of the original ancestry they are assigned by the prediction model and is significantly less computationally intensive than running software such as ADMIXTURE^8^ in the GenoTools pipeline.

Docker containers have been created to ensure ease of use that houses the proper dependencies to run the serialized models. The DockerHub link can be found in the Software Availability section below. GenoTools provides functionality to pull these Docker containers for use in the pipeline, both locally and on high-performance computing systems with Singularity (https://sylabs.io/singularity/).

We developed and trained distinct models tailored to two separate genotyping arrays, the NeuroBooster Array (NBA)^9^ and the NeuroChip Array (NCA)^10^. We focused on the overlap between each array and our reference panel. Both models are also appropriate for sequencing where there is a significant overlap of high-quality variants.

### Reference Panel

The reference panel for ancestry prediction models consists of 4008 samples from the 1000 Genomes Project (1000 Genomes)^11^, the Human Genome Diversity Project (HGDP)^12^, and an Ashkenazi Jewish reference panel^13^. The full counts per ancestry and quality control steps are detailed in **Figures 3 and 4** below.

**Figure 3.**
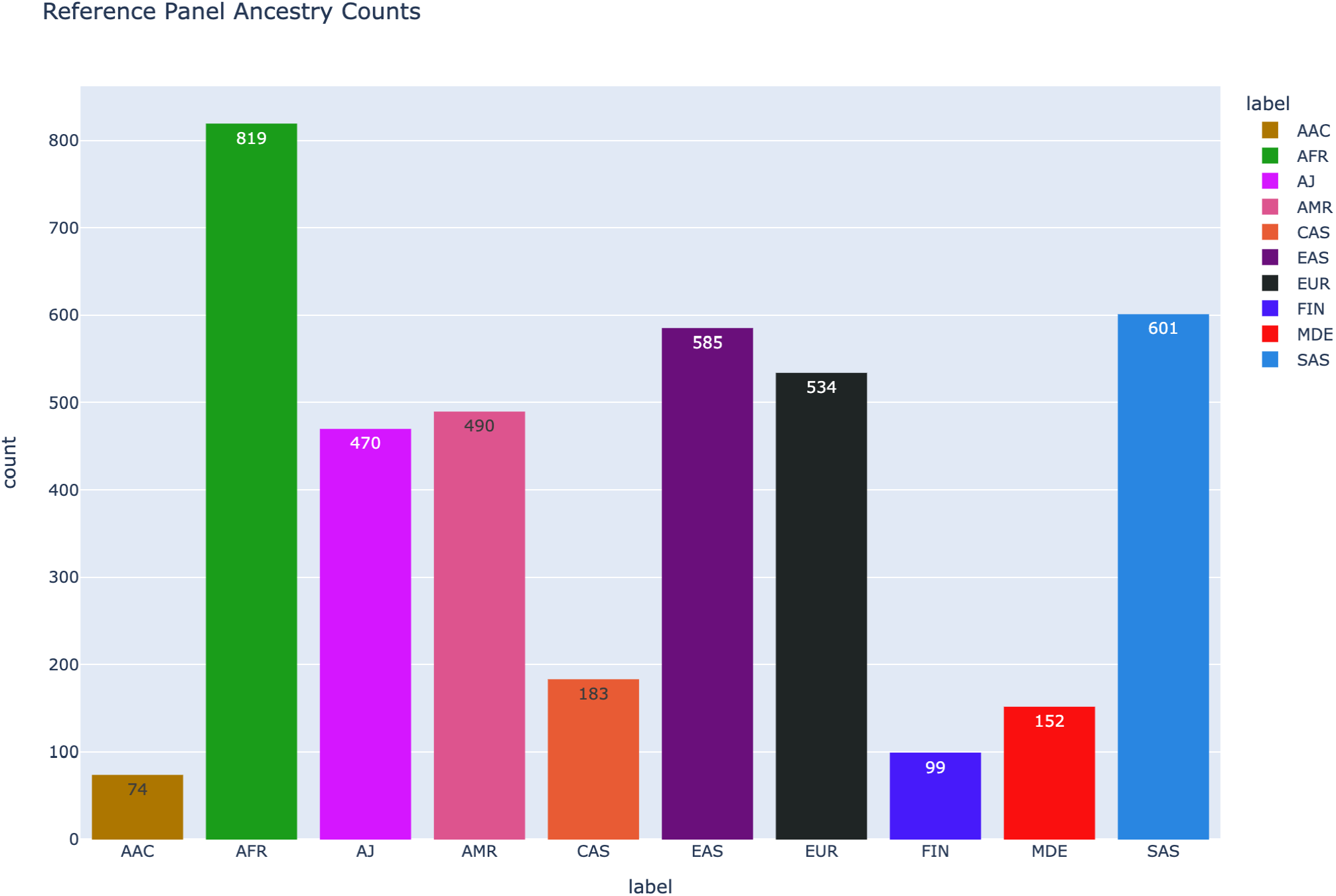
Reference panel ancestry counts.

**Figure 4.**
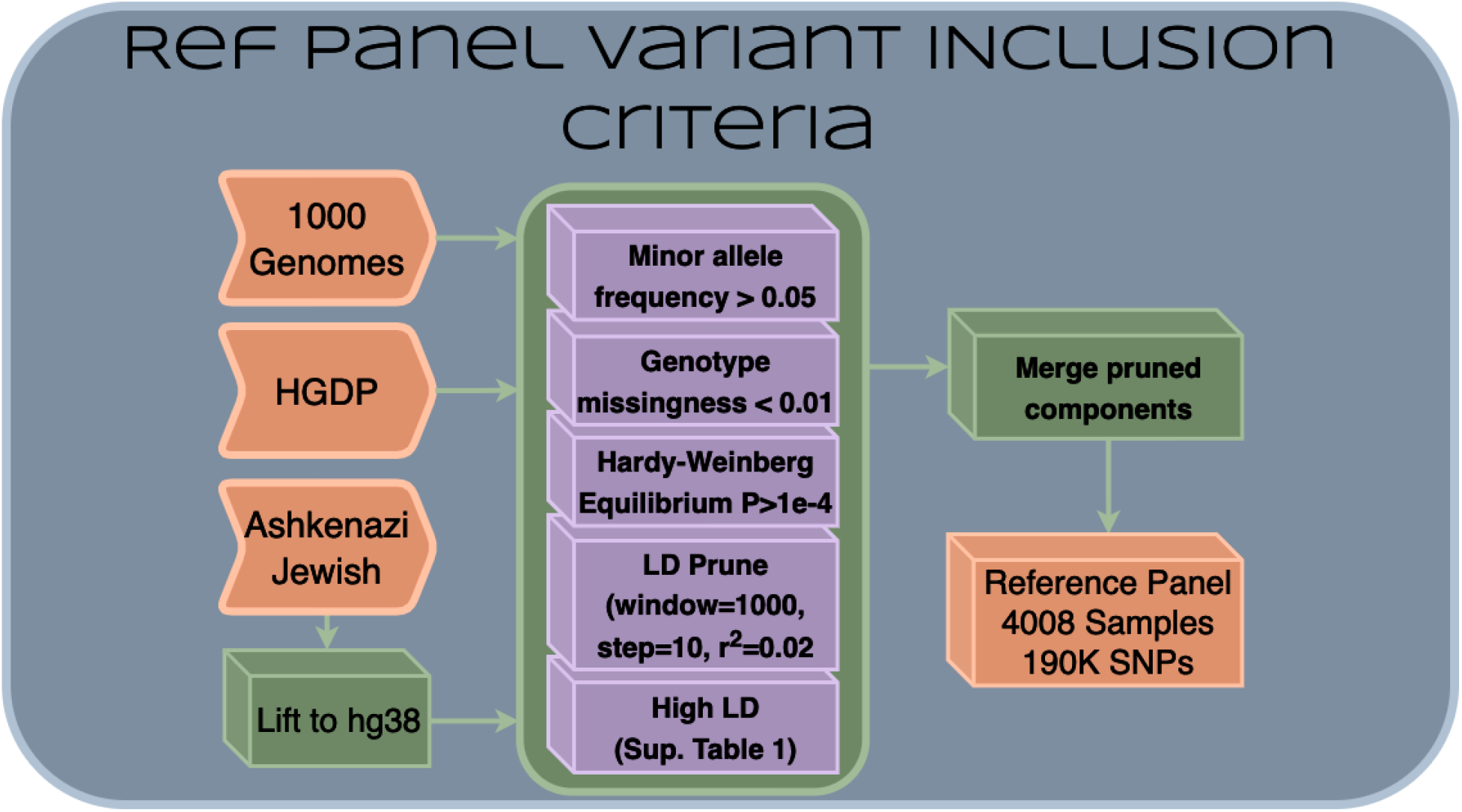
Reference panel variant inclusion criteria.

The AFR and AAC reference ancestry groups both come from the 1000 Genomes Project data and initially, their counts were 703 and 190 samples, respectively. We found the decision boundary formed by these initial labels heavily favored the prediction of AAC labels on new genotypes. To adjust this, a perceptron model (sklearn SVC with sigmoid kernel) was fit on the reference PCs for the AFR and AAC ancestry groups, to mathematically formalize a more appropriate decision boundary for ancestry prediction purposes. The reference labels were reassigned based on these predictions, which is reflected in the ancestry counts listed above.

Variants with a minor allele frequency less than 0.05 (--maf 0.05), with a missingness rate greater than 0.01 (--geno 0.01), or that deviated significantly from HWE (p < 1e-4) are removed from the reference panel. In addition, the reference panel has been excluded for LD (window size=1000, step size=10, r2=0.02) and only autosomal SNPs are included. Areas of high LD are also removed from the reference samples (**Table 1)**. Please note that this exclusion was performed independently on each set of samples (1000 Genomes, HGDP, and AJ) before merging to form the full reference panel (**Figure 4**).

**Table 1.** High LD Exclusion Regions (hg38)

| Chromosome | Start | End |
| --- | --- | --- |
| 2 | 94531009 | 94531062 |
| 6 | 24999772 | 33532223 |
| 6 | 26726947 | 26726981 |
| 8 | 8142478 | 12142491 |
| 11 | 50809938 | 50809993 |
| 11 | 54525074 | 55026708 |
| 11 | 55104000 | 55104074 |
| 12 | 34504916 | 34504974 |
| 20 | 28534739 | 28534795 |
| **Table 1.** High LD Exclusion Regions (hg38) |  |  |

### Quality Control

Following ancestry prediction, GenoTools applies stringent QC procedures at both the sample and variant levels to ensure data quality and eliminate potential biases or artifacts. The following quality control steps were developed for the Global Parkinson’s Genetics Program (GP2), acting as default parameters and thresholds for GenoTools.

### Sample-Level QC

1. Call rate exclusion, Default: 0.05.
2. Sex chromosome heterozygosity exclusion, Default: [0.25, 0.75] (F coefficient [female, male]).
3. Kinship, Defaults: 0.0884/0.354 related/duplicated:
4. Heterozygosity F, Default: [-0.15, 0.15]:

### Variant-Level QC

1. Missing Haplotype, Default: 1e-4 or 0.05/n for n>10,000
2. Hardy-Weinberg Equilibrium, Default: 1e-4, controls-only
3. Genotype Missingness, Default 0.05

### Association

#### Principal Component Analysis

1. Minor Allele Frequency (maf): Default: 0.01
2. Genotype Missingness (geno): Default: 0.01
3. Hardy-Weinberg Equilibrium (hwe). Default: 5e-6
4. Pairwise exclusion (indep_pairwise). Default: [1000, 10, 0.02] [window, step, variance inflation factor (VIF)]
5. Exclusion Regions: Based on the genome build (hg19/hg38), Hardcoded, high-LD regions.

#### General Linear Model

PLINK2 –glm runs an association on case/control or continuous phenotypes. Genomic inflation factor (λ) is calculated using the non-central chi-squared distribution (ncx2) function from the scipy library.

#### Outputs

GenoTools returns post-QC PLINK2 format genotype files divided by ancestry with pairwise relatedness information and summary statistics within each ancestry, and a JSON containing information following each step in the pipeline.

The JSON includes predicted counts per ancestry, predicted labels per sample, confusion matrix, balanced accuracy, PCs, and UMAP components from ancestry. Each QC step returns metrics, step name, whether the step passed or failed (True/False), whether the step was sample-level or variant-level, the ancestry group the step was run in, and the number and lists of IDs of samples or variants excluded at that step. For the association, a table containing the λ/λ_1000_ for each ancestry group that a GWAS was run on is output to the JSON (**Table 2**).

**Table 2.**
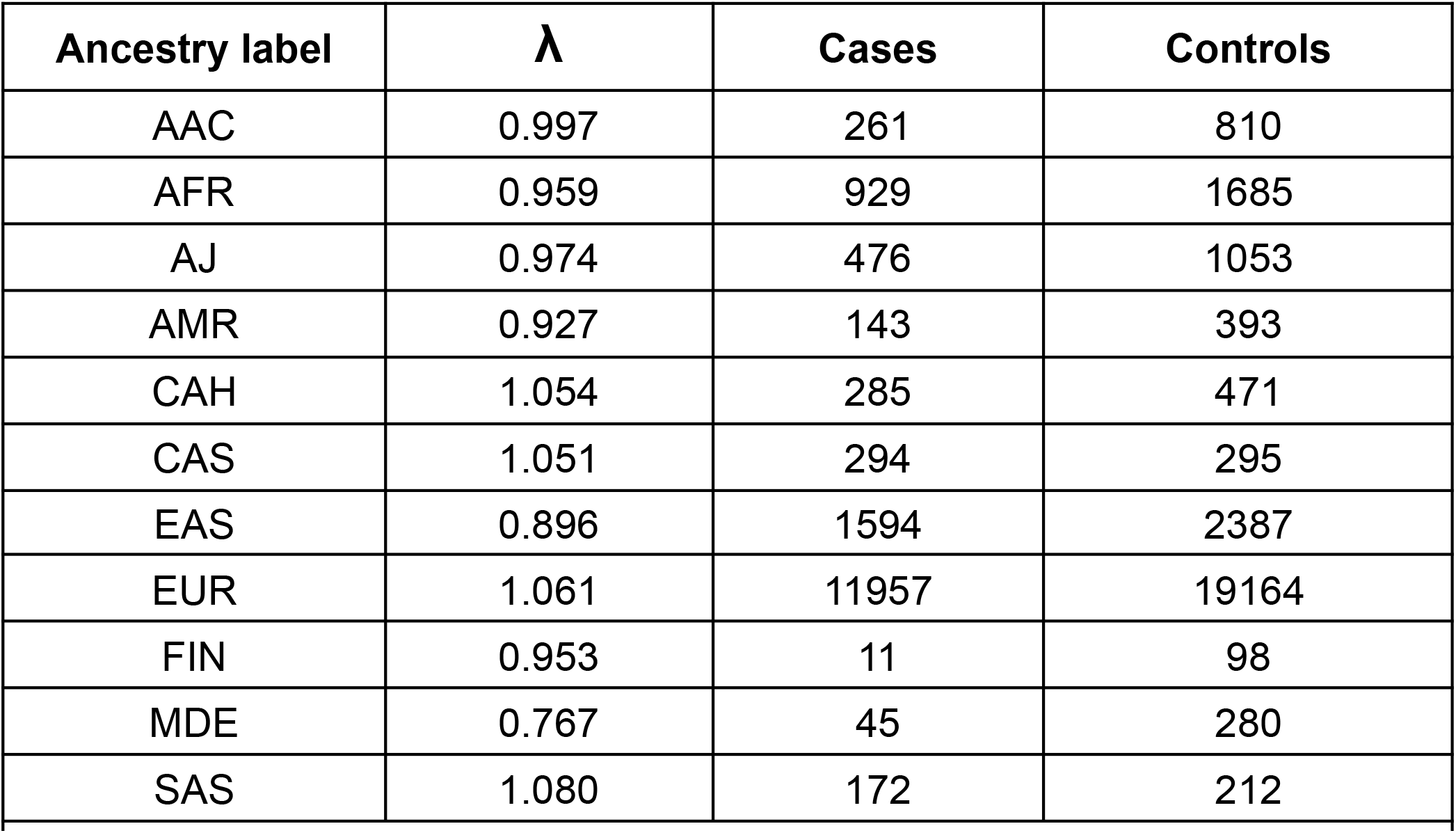
Genomic Inflation per-ancestry. EAS λ_1000_= 0.945, EUR λ_1000_= 1.004, and AFR λ_1000_= 0.966. λ_1000_ is used for case/control counts >1000. EUR (Europe), AJ (Ashkenazi Jewish), EAS (East Asian), SAS (South-Asian), CAH (complex admixture history), FIN (Finnish), AMR (Latino/Admixed American), MDE (Middle East), CAS (Central-Asian), AFR (African), AAC (African-Admixed).

## Results

GenoTools serves as the primary genotype processing pipeline of several large initiatives, including NIH’s Center for Alzheimer’s and Related Dementias (CARD) and the Global Parkinson’s Genetics Program (GP2)^14^. To date, GP2 has released 47,853 samples across 11 predicted ancestries. Benchmarking was performed on 10,000 samples with 1,954,758 variants with the following specifications: 4 hours, 35.5 minutes with 15.3 GB memory for the full pipeline with training, and 49.5 minutes with 20.3 GB memory for the full pipeline using a pre-trained model. In GP2’s Release 6^15,^ containing 44,831 samples, GenoTools was used for ancestry and quality control achieving a 95.9 +/- 1.4% balanced accuracy, with 0.969 precision and 0.964 F1 score with training and validation on the reference panel; See the confusion matrix in **Figure 5**.

**Figure 5.**
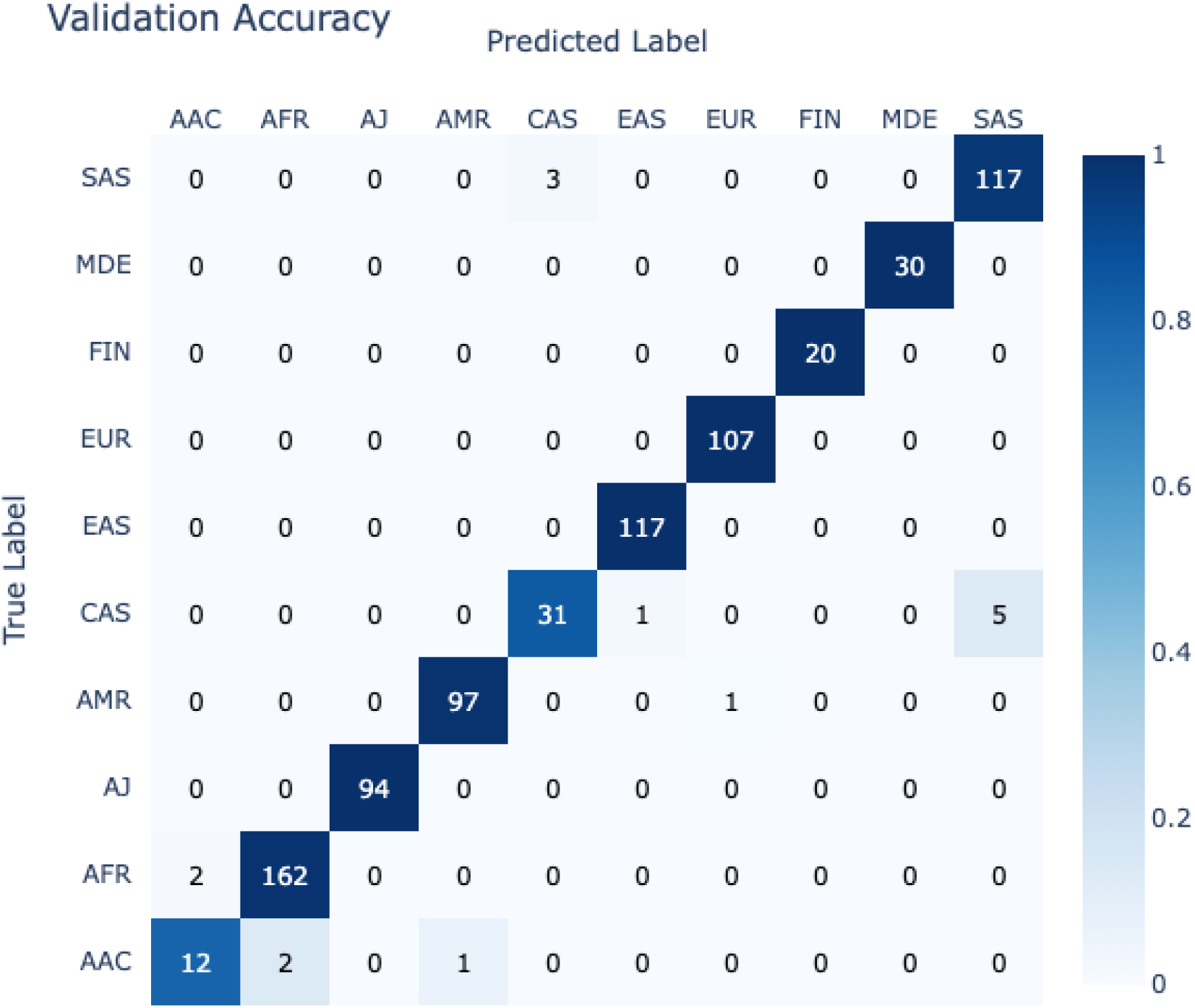
Ancestry model validation accuracy by ancestry label.

To show how GenoTools can be used at scale, the standard pipeline without the relatedness check was run on genotypes in the UK Biobank^16^. Benchmarking was performed on 487,158 samples with 1,033,310 variants in pgen format with the following specifications: 16 hours, 48.5 minutes with 502 GB memory for the full pipeline with training, and 8 hours, 51 minutes with 502 GB memory for the full pipeline using a pre-trained model. Kinship estimation for sample sizes over 50,000 is a bottleneck in the pipeline. To demonstrate this, when relatedness pruning was run on 100,000 predicted EUR ancestry samples in the UK Biobank data it performed with the following specifications: 39 hours, 16.5 minutes with 94.6 GB memory. Full PCs and CAH-delineated PCs can be seen in **Figures 6 and 7** below.

**Figure 6.**
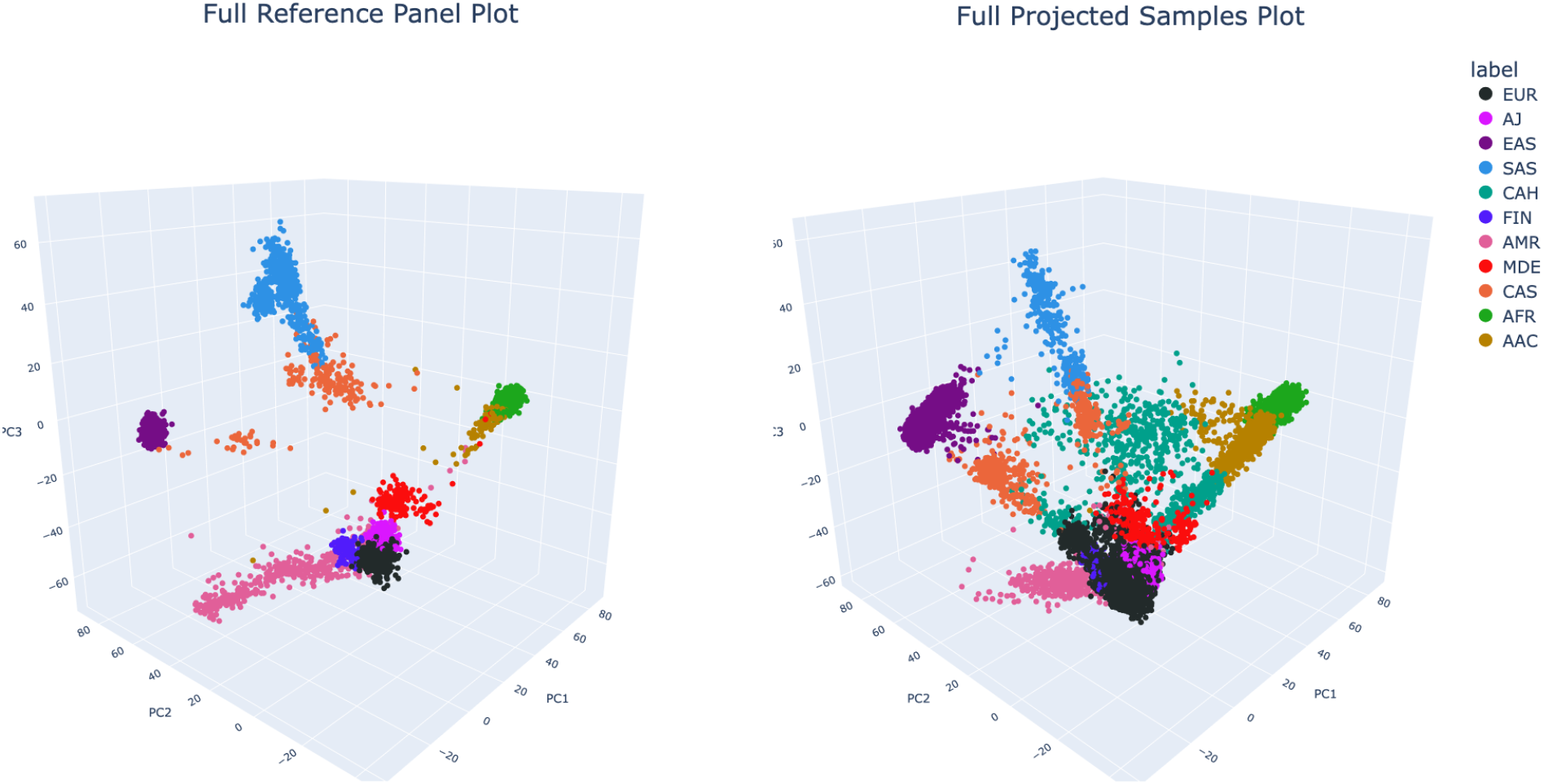
Reference Panel (left) and GP2 Release 6 samples (right). Each point represents a sample and the colors depict the ancestral background as shown in the color legend: EUR = black, AJ = magenta, EAS = purple, SAS = blue, CAH = light green, FIN = blue, AMR = pink, MDE = red, CAS = orange, AFR = green, AAC = yellow. EUR (Europe), AJ (Ashkenazi Jewish), EAS (East Asian), SAS (South-Asian), CAH (complex admixture history), FIN (Finnish), AMR (Latino/Admixed American), MDE (Middle East), CAS (Central-Asian), AFR (African), AAC (African-Admixed).

**Figure 7.**
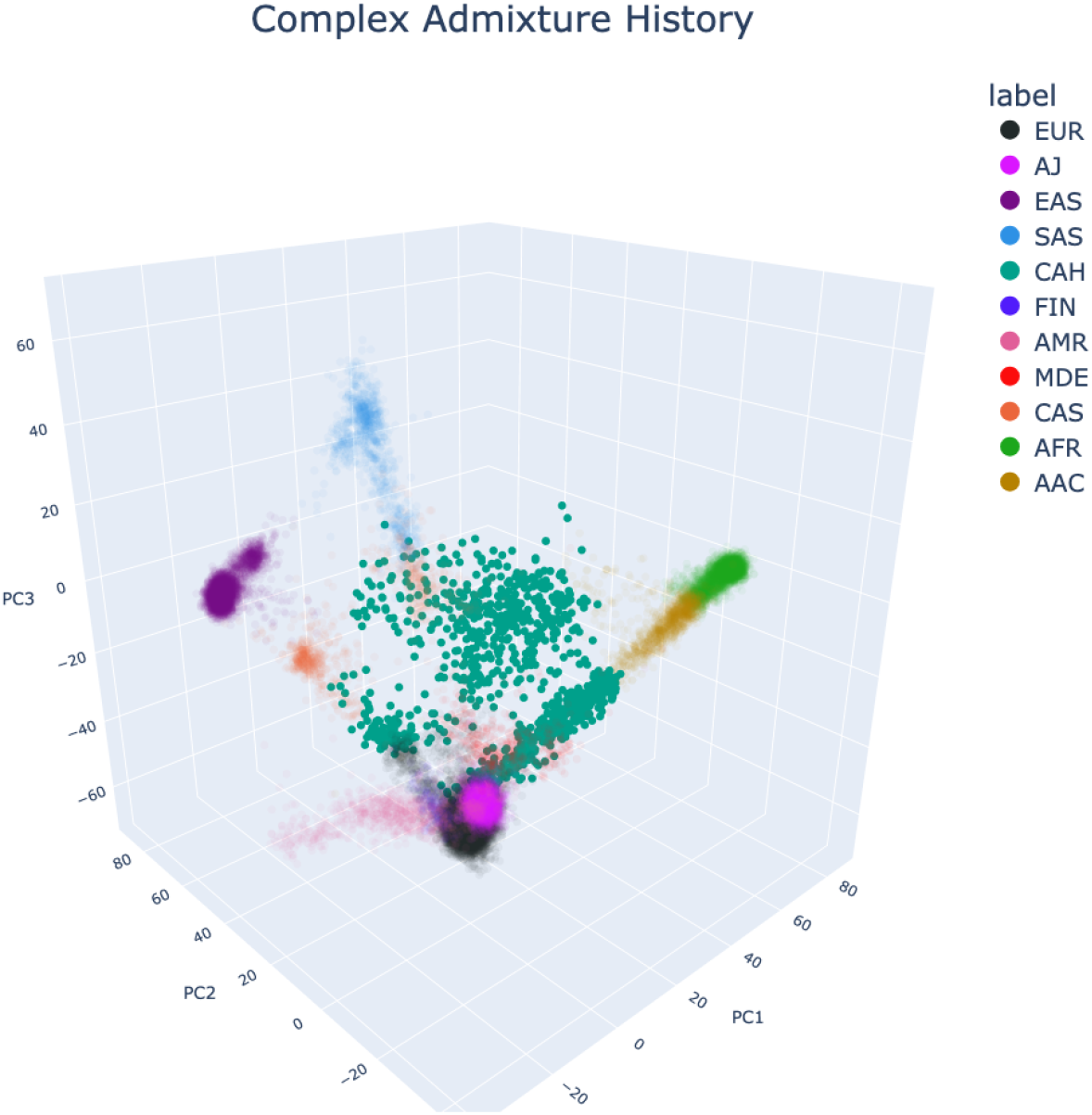
Complex admixture history (CAH) PCs. Each point represents a sample and the colors depict the ancestral background as shown in the color legend: EUR = black, AJ = magenta, EAS = purple, SAS = blue, CAH = light green, FIN = blue, AMR = pink, MDE = red, CAS = orange, AFR = green, AAC = yellow. EUR (Europe), AJ (Ashkenazi Jewish), EAS (East Asian), SAS (South-Asian), CAH (complex admixture history), FIN (Finnish), AMR (Latino/Admixed American), MDE (Middle East), CAS (Central-Asian), AFR (African), AAC (African-Admixed).

Additionally, we found a 66249.598 chi-square, p-value <1e-16, with 50 degrees of freedom, and 0.654 Cramér’s V between predicted ancestry labels and self-reported race information in GP2 (Figure 8). In the UK Biobank^15^ data, we found a 1526563.448 chi-square, <1e-16 p-value, 180 degrees of freedom, and 0.564 Cramér’s V between predicted ancestry and self-reported race (Figure 9), suggesting strong relationships between predictions and self-report in both datasets.

**Figure 8.**
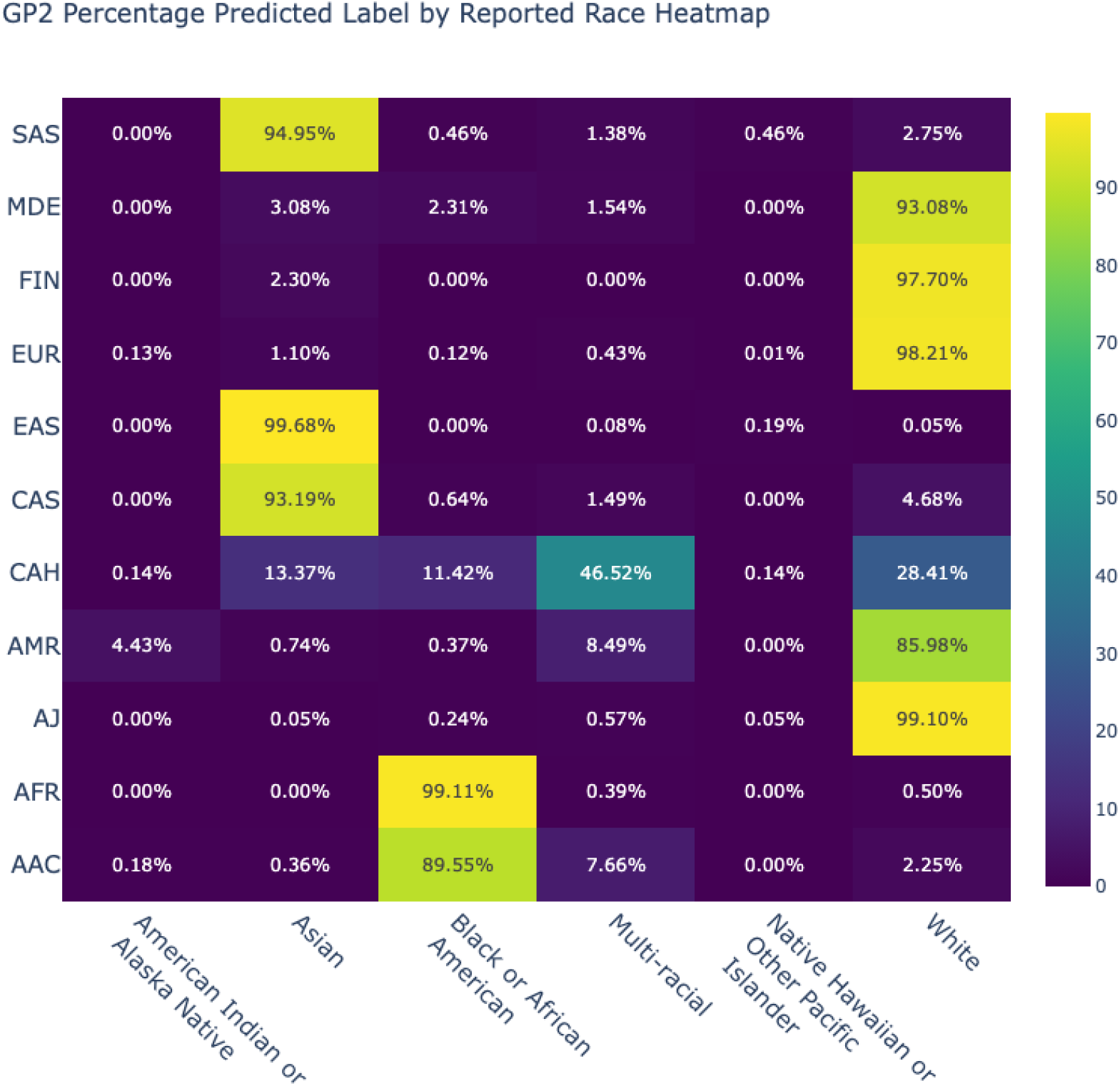
Comparison of predicted genetic ancestry and self-reported race for GP2 Release 6.

**Figure 9.**
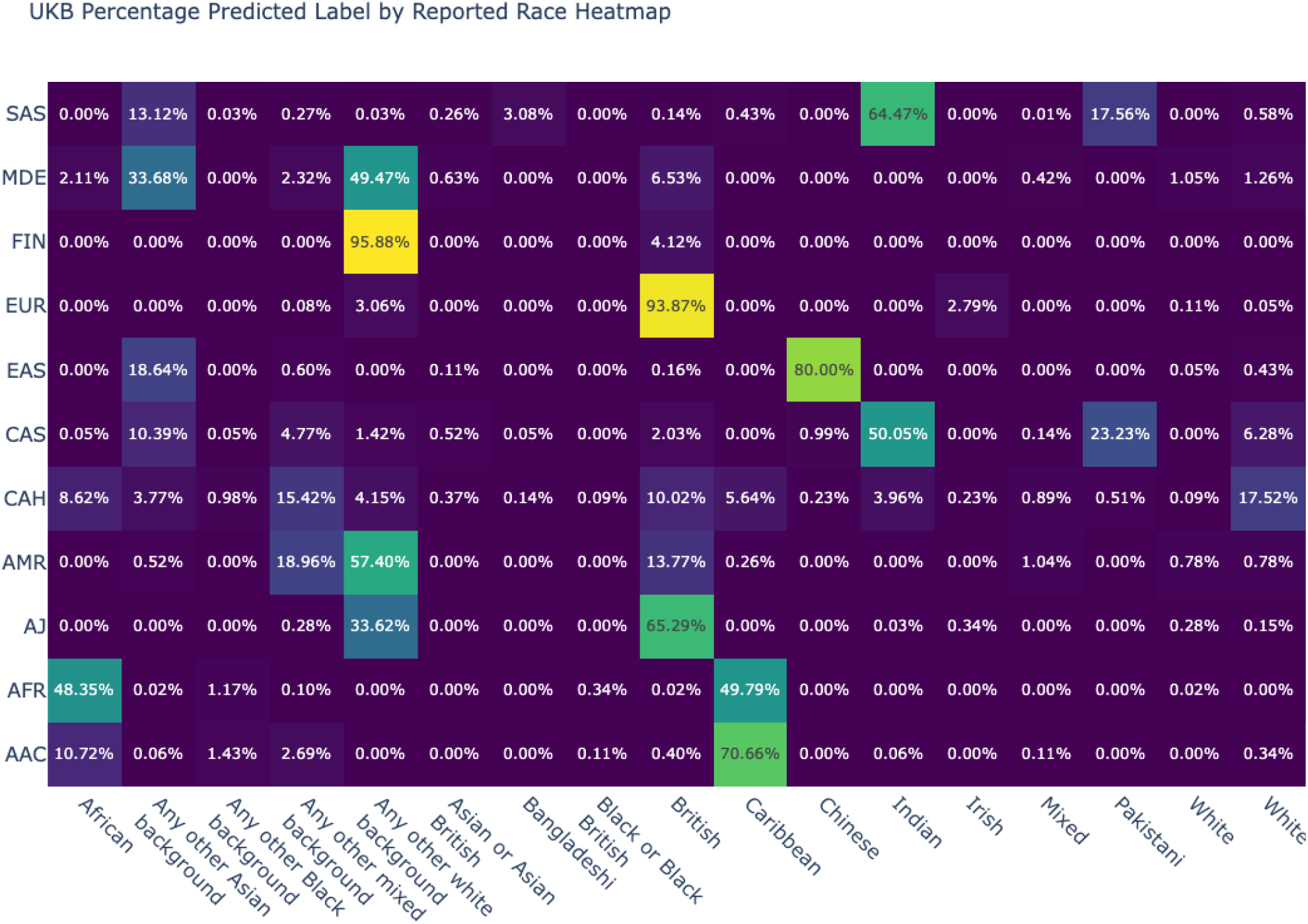
Comparison of predicted genetic ancestry and self-reported race for UK Biobank

**Figure 10.**
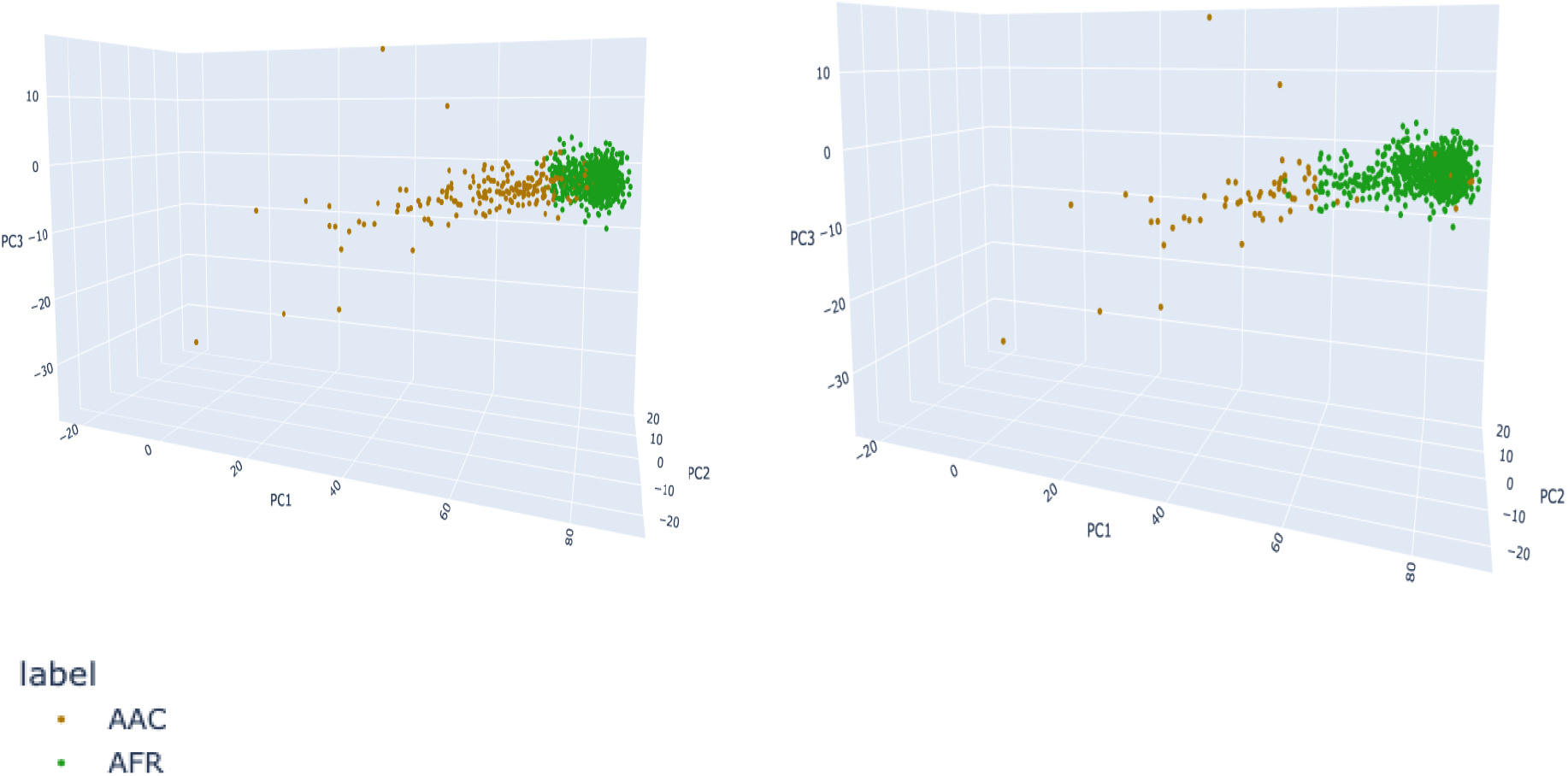
1000 Genomes AFR and AAC PCs (left) and Perceptron-labeled AFR and AAC PCs (right)

GenoTools QC on GP2 Release 6 excluded 2,414 samples across 11 assigned ancestry labels yielding genomic inflations indicating reliable data across all ancestries (Table 1). GP2 Release 6 demonstrated GenoTools’ flexibility by simultaneously processing samples across multiple localities in both Google Cloud infrastructure and local clusters, across varying compute solutions (SLURM, Google Batch, etc), with a single command, within a few hours of runtime and minimal compute.

The development of the GP2 dataset with GenoTools has led to published novel results in diverse populations, as demonstrated in a study of Parkinson’s Disease (PD) in African and African Admixed populations, which identified novel common risk factors for PD in the *GBA1* locus that suggests decreased glucocerebrosidase activity^17^. In this study, 197,918 total samples (1,488 cases and 196,430 controls), including GP2 data, were processed by GenoTools. This demonstrates the power and flexibility of the methods to rapidly process multiple datasets as well as generate accurate ancestry predictions and top-quality data at scale to streamline ancestry-specific studies.

GenoTools has also been used for QC and ancestry prediction in cell lines of 225 patients for a study into pathogenic variants in *GBA1* in Gaucher disease (GD) type1 (GD1) for risk of PD in GD1 patients^18^, further demonstrating the flexibility of GenoTools to studies of all sizes, across different genetics datasets.

## Discussion

GenoTools has facilitated the development and scaling of a large PD genetics dataset in GP2 and generated novel results in ancestrally diverse population genetics data. It has been used to streamline and automate the data processing and sample management involved in a large-scale, multi-national, asynchronous genetics program, processing data in multiple localities around the world in parallel, allowing for harmonization across studies and data silos. This Python package has taken the standard workflows of population genetics and has distilled down a process requiring many complex and time-consuming steps into single-command workflows that are fully customizable to the user’s specifications and can be run anywhere. GenoTools provides efficient and robust ancestry prediction, quality control, and summary statistics for genotyping array, WGS, and exome data, and ensures that researchers can trust their results across diverse populations and reach results with little effort for replicable, open science.

## Software Availability

- GitHub: GenoTools v1.2.1 https://github.com/dvitale199/GenoTools
- Pypi: https://pypi.org/project/the-real-genotools/
- Zenodo: https://zenodo.org/doi/10.5281/zenodo.10443257
- Reference panel: https://storage.googleapis.com/genotools_refs/ref_panel/1kg_30x_hgdp_ashk_ref_panel.zip
- Models
  - NeuroBooster: https://storage.googleapis.com/genotools_refs/models/nba_v1.zip
  - NeuroChip: https://storage.googleapis.com/genotools_refs/models/neurochip_v1.zip
  - WGS (all 190k reference panel SNPs): https://storage.googleapis.com/genotools_refs/models/wgs_v1.zip
- Dockerhub: https://hub.docker.com/repository/docker/mkoretsky1/genotools_ancestry/general
- Licensing Information: Apache License, Version 2.0, January 2004 (http://www.apache.org/licenses/)

## Acknowledgments, Funding and COI

This research was supported in part by the Intramural Research Program of the NIH, National Institute on Aging (NIA), National Institutes of Health, Department of Health and Human Services; project number ZO1 AG000534, as well as the National Institute of Neurological Disorders and Stroke.

Data used in the preparation of this article were obtained from Global Parkinson’s Genetics Program (GP2). GP2 is funded by the Aligning Science Across Parkinson’s (ASAP) initiative and implemented by The Michael J. Fox Foundation for Parkinson’s Research (**https://gp2.org**). For a complete list of GP2 members see **https://gp2.org**.

D.V., H.L.L., H.I., F.F., K.L., and M.A.N. participated in this project in a competitive contract awarded to DataTecnica LLC by the National Institutes of Health to support open science research. M.A.N. also currently serves on the scientific advisory board for Character Bio Inc., is a scientific founder at Neuron23 Inc., and owns stock.

## Notes

https://github.com/dvitale199/GenoTools

https://pypi.org/project/the-real-genotools/

https://storage.googleapis.com/genotools_refs/ref_panel/1kg_30x_hgdp_ashk_ref_panel.zip

https://storage.googleapis.com/genotools_refs/models/nba_v1.zip

https://storage.googleapis.com/genotools_refs/models/neurochip_v1.zip

https://storage.googleapis.com/genotools_refs/models/wgs_v1.zip

https://hub.docker.com/repository/docker/mkoretsky1/genotools_ancestry/general

https://zenodo.org/doi/10.5281/zenodo.10443257

### Competing Interest Statement

D.V., H.L.L., H.I., F.F., K.L., and M.A.N.'s participation in this project was part of a competitive contract awarded to DataTecnica LLC by the National Institutes of Health to support open science research. M.A.N. also currently serves on the scientific advisory board for Character Bio Inc plus is a scientific founder at Neuron23 Inc and owns stock.

### Summary of Updates

adding a missing citation and minor grammar edits

